# Natural selection contributed to immunological differences between human hunter-gatherers and agriculturalists

**DOI:** 10.1101/487207

**Authors:** Genelle F Harrison, Joaquin Sanz, Jonathan Boulais, Michael J Mina, Jean-Christophe Grenier, Yumei Leng, Anne Dumaine, Vania Yotova, Christina M. Bergey, Stephen J. Elledge, Erwin Schurr, Lluis Quintana-Murci, George H. Perry, Luis B. Barreiro

**Affiliations:** Department of Human Genetics, Faculty of Medicine, McGill University, Montreal, QC, Canada.; Department of Genetics, CHU Sainte-Justine Research Center, Montreal, QC, Canada.; Department of Biochemistry, Faculty of Medicine, Université de Montréal, QC, Canada.; Department of Pathology, Brigham and Women’s Hospital, Harvard Medical School, 75 Francis Street, Boston, MA 02115, United States of America.; Department of Genetics, Harvard Medical School and Division of Genetics, Brigham and Women’s Hospital, Howard Hughes Medical Institute, 77 Avenue Louis Pasteur, Boston, MA 02115, United States of America.; Departments of Anthropology and Biology, Pennsylvania State University, University Park, PA, USA.; Unit of Human Evolutionary Genetics, Institut Pasteur, Paris, France.; Centre National de la Recherche Scientifique, UMR2000, Paris, France.; Center of Bioinformatics, Biostatistics and Integrative Biology, Institut Pasteur, Paris, France.; Huck Institutes of the Life Sciences, Pennsylvania State University, University Park, PA, USA; University of Montreal, Department of Pediatrics, Montreal, QC, Canada.; University of Chicago, Department of Medicine, Section of Genetic Medicine, Chicago, USA.

## Abstract

The shift from a hunter-gatherer (HG) to an agricultural (AG) mode of subsistence is believed to have been associated with profound changes in the burden and diversity of pathogens across human populations. Yet, the extent to which the advent of agriculture may have impacted the evolution of the human immune system remains unknown. Here we present a comparative study of variation in the transcriptional responses of peripheral blood mononuclear cells to bacterial and viral stimuli between Batwa rainforest hunter-gatherers and Bakiga agriculturalists from Uganda. We observed increased divergence between hunter-gatherers and agriculturalists in the transcriptional response to viruses compared to that for bacterial stimuli. We demonstrate that a significant fraction of these transcriptional differences are under genetic control, and we show that positive natural selection has helped to shape population differences in immune regulation. Across the set of genetic variants underlying inter-population immune response differences, however, the signatures of positive selection were disproportionately observed in the rainforest hunter-gatherers. This result is counter to expectations based on the popularized notion that shifts in pathogen exposure due to the advent of agriculture imposed radically heightened selective pressures in agriculturalist populations.

---

The agricultural revolution beginning 10,000-12,000 BP was associated with profound changes in human ecology^1^, which in turn are hypothesized to have precipitated major new infectious disease burdens^2-4^. Specifically, the construction of permanent settlements and a subsequent increase in population density associated with the agricultural transition^5,6^ may have facilitated the establishment and transmission of infectious agents such as smallpox, measles, rubella, and other pathogens that require hundreds to thousands of host individuals to spread and persist^7,8^. Agriculturalists and pastoralists also lived in proximity with their domesticated animals, providing opportunity for novel or expanded zoonotic transmission^4^ of pathogens potentially including rotavirus, measles virus, and influenza^9-11^. Finally, agriculturists performed extensive modifications to the landscape, including clearing fields and constructing irrigation systems, which may have led to an increase in the incidence of vector-borne diseases, such as *Plasmodium falciparum* malaria^12,13^.

Consequentially, the transition to an agriculturalist lifestyle is hypothesized to have contributed to the strong genetic signatures of recent positive selection that are repeatedly observed within or nearby immune-related genes in worldwide agriculturalist populations^14,15^. However, the absence of comparative functional studies from pairs of populations that differ in their modes of subsistence, i.e., hunter-gatherers (HG) versus agriculturalists (AG), have thus far precluded the development of hypotheses concerning specifically how the agricultural transition may have impacted evolution of human immune system diversity. To begin studying this topic, we used a combination of evolutionary genomic and functional immunological tools to study differences in immune responses between the Batwa, a rainforest hunter-gatherer population from southwest Uganda, and their Bantu-speaking agriculturalist neighbors, the Bakiga.

## Results

### Significant Batwa-Bakiga immune response differences

Whole blood samples from 103 individuals (59 HG-Batwa and 44 AG-Bakiga, Supplementary Figure 1) were collected, and peripheral blood mononuclear cells (PBMCs) from these samples were isolated and cryopreserved. PBMCs were collected and processed for both populations simultaneously during the same field expedition to minimize technical variability. Each individual was genotyped for ~1 million genome wide SNPs^16^, with additional imputation to 10,530,212 SNP genotypes (see Materials and Methods). These data were used to estimate genome-wide levels of HG-Batwa and AG-Bakiga ancestry, using the program ADMIXTURE^17^. We observed variable but considerable levels of AG-Bakiga ancestry among self-identified HG-Batwa individuals (mean = 21.0 %; range = 0 – 93.3%). However, estimated levels of HG-Batwa ancestry among self-identified AG-Bakiga individuals were typically lower (mean = 4.3%; range = 0 – 9.7%, Figure 1A).

**Fig. 1.**
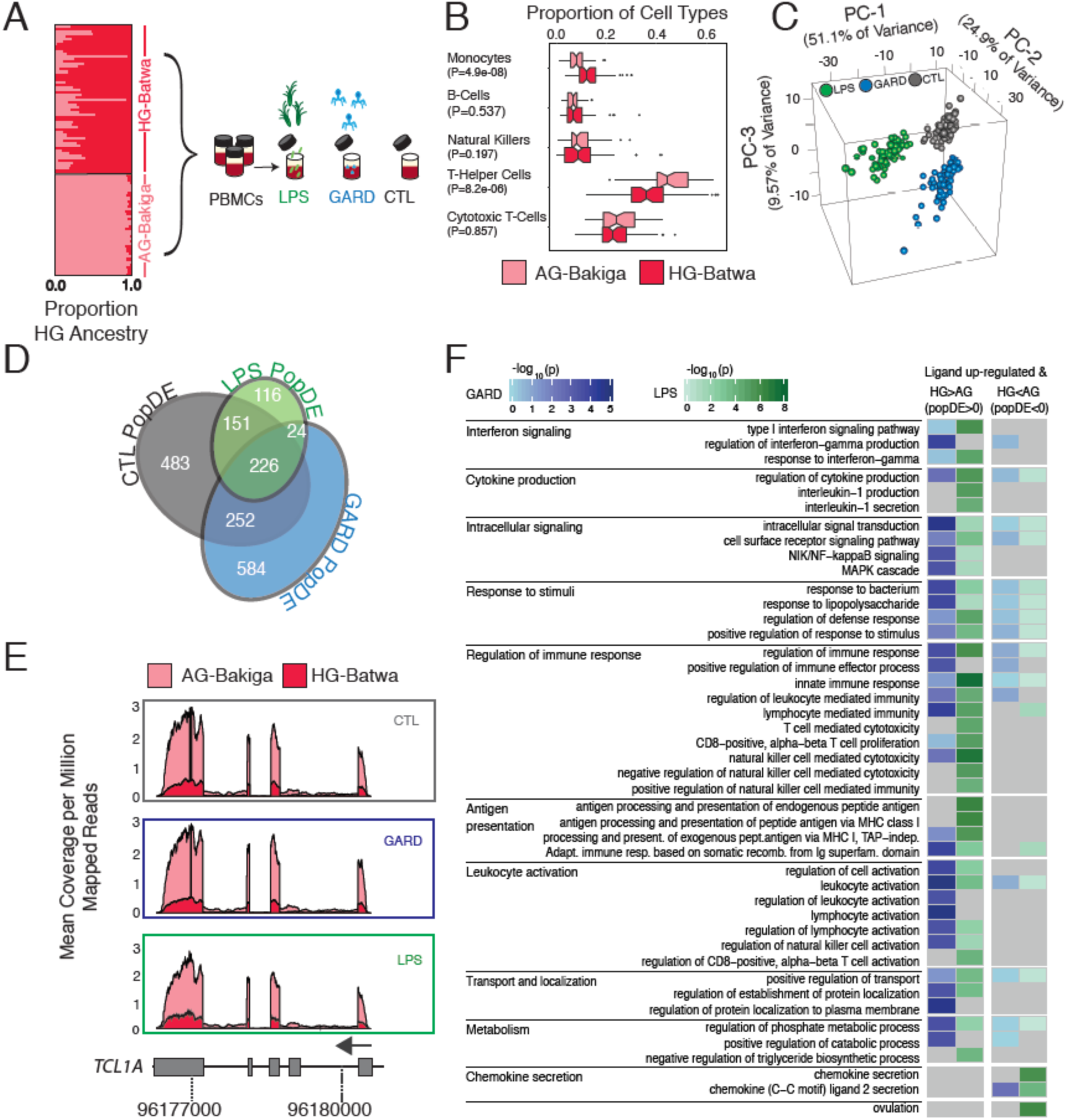
Transcriptional differences between Batwa hunter-gatherer and Bakiga agriculturalist populations. (**A**) Schematics of the study design. The structure plot to the left shows the proportion of HG-ancestry (dark pink) and AG-ancestry (light pink) for each individual included in the study. Their placement along the Y-axis corresponds to how they self-identified. (**B**) Boxplots of the proportions of the main cell types found in PBMCs in the Batwa (dark pink) and the Bakiga (light pink). (**C**) Principal components analysis of gene-expression data. The first three PCs separate non-infected PBMCs from PBMCs stimulated with either LPS or GARD. (**D**) Venn diagram of PopDE genes detected in each condition. (**E**) Example of a PopDE gene (*TCL1A*) in which gene expression is higher in the AG population (light pink) than the HG population (dark pink) in all conditions. Expression is shown as the mean coverage per genomic position (corrected by total mapped reads) per individual in each population. **F)** Gene Ontology terms enriched among PopDE genes in the GARD (blue) and LPS (green) conditions. The left and right columns illustrate the level of significance of the enrichments (-log10 p values) for genes showing higher gene expression levels in HG and AG populations, respectively. For visual purposes, we only show a subset of all significant GO terms (see Materials and Methods). A full list is provided in Supplementary Table 3.

To characterize variation in the immune response between HG-Batwa and AG-Bakiga populations we exposed PBMCs to Gardiquimod (GARD, TLR7 agonist), which mimics an infection with a single-stranded RNA virus, and lipopolysaccharide (LPS, TLR4 agonist), which simulates an infection with gram-negative bacteria. We also maintained an unexposed control in the same experimental conditions (CTL). Following 4 hours of stimulation, we collected RNA-sequencing data from matched non-stimulated and stimulated PBMCs (Figure 1A). Following quality control filtering we analyzed high-quality RNA-sequencing profiles (n=229 RNA-sequencing profiles across treatment combinations) from 99 individuals (57 HG-Batwa and 42 AG-Bakiga; see Methods, Supplementary Figure 1, and Supplementary Table 1). To confirm successful ligand stimulation, we performed a principal component analysis (PCA) on the correlation matrix of normalized gene expression levels for all conditions. The first PC explained 51.1% of the variance in the expression values, and effectively separated the LPS condition from an unstimulated control (CTL). The combination of the second and third PCs further separated the GARD-stimulated PBMCs from the CTL cells (Figure 1C). As expected, the set of genes up-regulated in response to both stimuli were significantly enriched (False Discovery Rate (FDR)<1×10^−15^) for genes known to be involved in immune defense and inflammatory responses, with a particularly strong enrichment for anti-viral response genes in the GARD condition (Supplementary Table S3).

Because PBMCs are a composite of various innate and adaptive immunity cell types, we first determined whether there were differences in the cellular compositions of PBMCs between the HG-Batwa and AG-Bakiga. Using fluorescence-activated cell sorting (FACS) we estimate the proportion of each of the major cell types comprising PBMCs for every individual (see Supplementary Figure 2 for a summary of the gating strategy). We found that the proportion of CD14^+^ monocytes was higher in individuals with greater HG-Batwa ancestry (*P* = 4.9×10^−08^), while the proportion of CD3^+^/CD4^+^ helper T-cells was higher in individuals with greater AG-Bakiga ancestry (*P* = 8.2×10^−06^; Figure 1B). Using linear models that account for variation in cell composition, sex, and additional technical covariates (see Materials and Methods), we next identified genes whose expression levels were linearly correlated with ancestry within each of the experimental conditions (i.e., population differentially expressed, or PopDE genes). Of the 10,885 expressed genes tested, 1,836 genes (16.9% of the total) were found to be PopDE (FDR < 0.05) in at least one condition (Figure 1D with 1E for an example). Among PopDE genes, genetic ancestry explains, on average, 14.4% (Quantile 5%-95% interval: 6.8-25.1) of the overall variance in gene expression observed among individuals, an amount comparable to the proportion of variation that can be attributed to differences in cell composition (mean = 16.8%; Quantile 5%-95% interval: 2.9-39.0) and much higher than the proportion explained by sex (mean = 3.4%; Quantile 5%-95% interval: 0.2-9.8; Supplementary Figure 3).

**Fig. 2.**
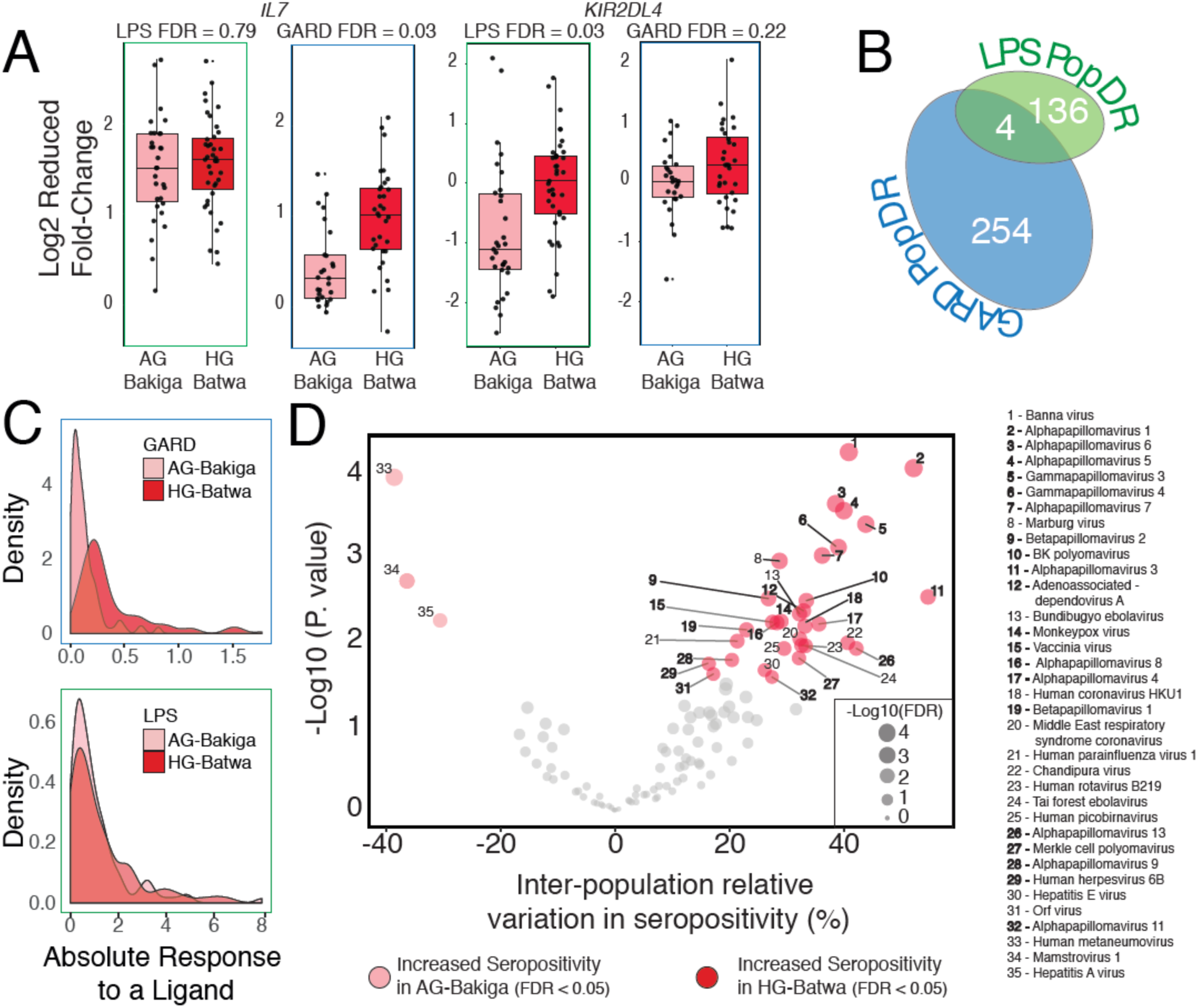
Differences in immune response between HG and AG populations. (**A**) Examples of two PopDR genes involved in immune response. The y axis shows the log2 fold changes in gene expression levels in response to LPS and GARD, for individuals from each of the two populations (x axis). (**B**) Venn diagram showing the number of PopDR genes identified in the LPS and GARD conditions. (**C**) Density plots showing the distributions of the absolute response to LPS and GARD of PopDR genes in each population. (**D**) A volcano plot showing an increase in seropositivity in the HG-Batwa population for 32 of the 130 viruses tested. Double stranded DNA-viruses showing a significant dependence to ancestry are marked in bold.

**Fig. 3.**
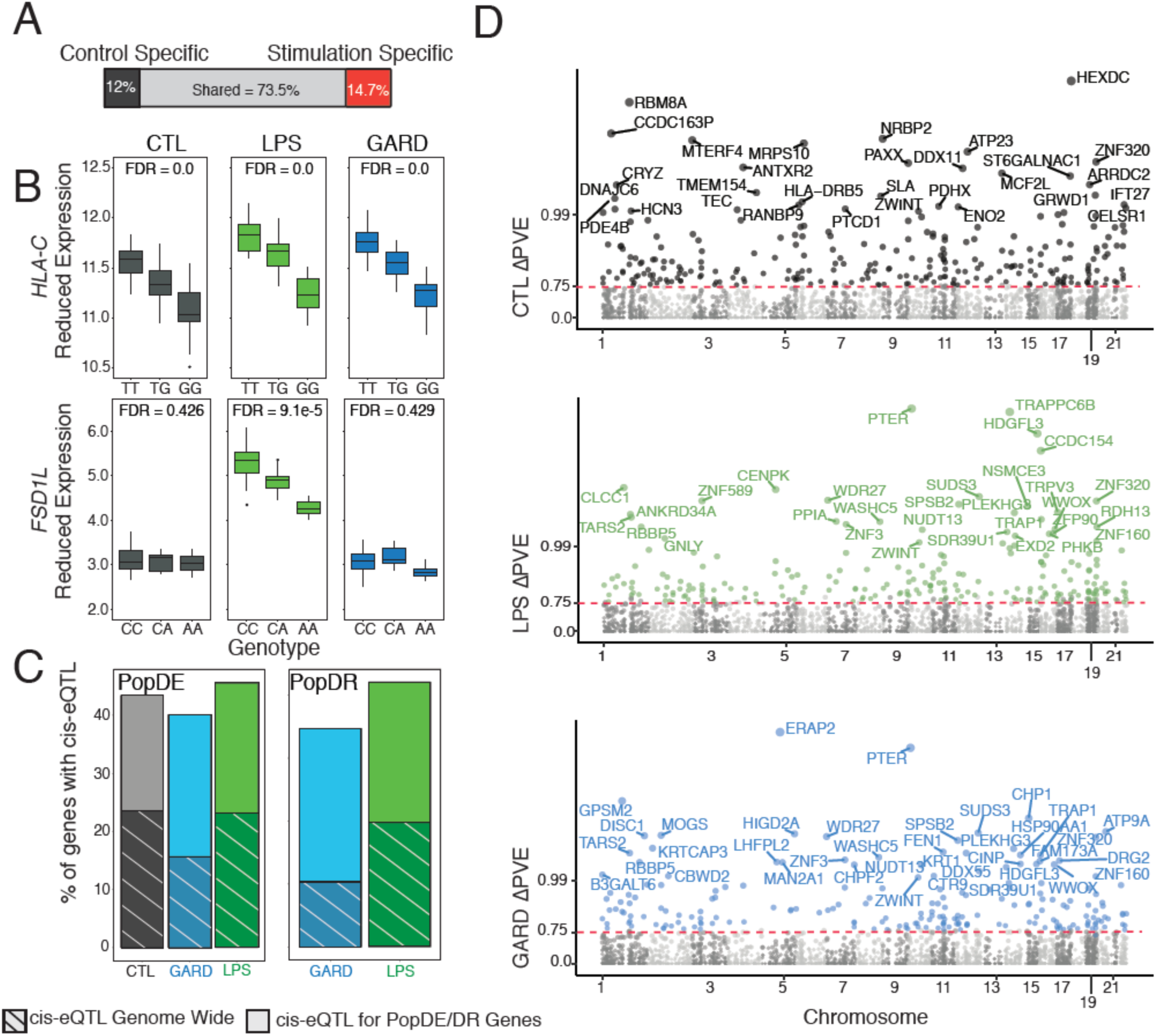
Analysis of the contribution of genetics to differences in immune response between the HG-Batwa and the AG-Bakiga. (**A**) Schematic representation of the number of cis-eQTL shared across all conditions, or only found in non-infected PBMCs, or found in LPS and/or GARD stimulated PBMCs (stimulation-specific eQTL). Stimulation-specific eQTL were defined as those showing very strong evidence of eQTL in the stimulated cells (FDR < 0.05), and very limited evidence in the non-infected cells (FDR always higher than 0.25). (**B**) Example of two cis-eQTL. The top example, *HLA-C*, was found across all experimental condition (CTL-FDR = 0.0, LPS-FDR = 0.0, GARD-FDR = 0.0). The bottom example, Fibronectin Type III and SPRY Domain Containing 1 Like (*FSD1L*) was detected exclusively in the LPS condition. In this example expression is in log2(counts per million) (CTL-FDR = 0.426, LPS-FDR = 9.09^−5^, GARD-FDR = 0.429). (**C**) Bar graphs showing an enrichment of genes containing cis-eQTLs among PopDE/PopDR genes (totality of bars) per compared to genome wide expectations (stripes). (**D**) Manhattan plot showing ∆PVE of cis-eQTL (normalized as -log10(1-∆PVE for easier viewing) on the Y-axis across all chromosomes for CTL (gray), GARD (blue), and LPS (green). Colored points have an FDR < 0.1 and a delta-PVE > 0.75. Points are labeled with the corresponding gene name when the PVE is > 0.99.

A Gene Ontology (GO) analysis did not reveal any particular biological pattern among genes showing higher expression levels in AG-Bakiga individuals (relative to HG-Batwa individuals) in the LPS- and GARD-stimulated PBMCs (Supplementary Table 3). In stark contrast, the set of genes with higher expression levels in HG-Batwa individuals following stimulation were markedly enriched in genes involved in relevant immunological processes such as interleukin-1 production (LPS: FDR = 2.5×10^−4^), regulation of interferon gamma production (GARD: FDR = 1.3×10^−2^) or leukocyte activation (GARD: FDR = 1.2×10^−2^; LPS: FDR = 5.7×10^−3^), among others (Figure 1F, Supplementary Table 3). These results suggest that increased HG-Batwa ancestry is associated with a generally stronger degree of immune activation.

### Viruses were likely the main driver of Batwa-Bakiga immune response differences

Several lines of evidence indicate that the regulation of the immune response to viral stimuli between HG-Batwa and the AG-Bakiga individuals is more divergent compared to that for bacterial stimuli. Among “stimuli-responsive genes” (i.e., the set of genes that exhibit expression changes upon LPS- or GARD-stimulation), we identified almost twice as many PopDE genes in the GARD condition as compared to the LPS condition (10.1% of all genes that respond to GARD *vs* 5.9% of all genes that respond to LPS; Chi-squared test, *P* < 2.2×10^−16^). When considering the set of genes for which the intensity of the response to LPS and GARD – defined as the fold-change in the stimulated condition relative to the unstimulated condition – varied as a function of genetic ancestry (i.e., population differentially responsive, or PopDR genes, Figure 2A for an example), we again observed approximately twice as many PopDR genes (FDR < 0.1) in GARD-stimulated cells compared to LPS-stimulated cells (258 PopDR for GARD vs. 140 PopDR for LPS, Figure 2B).

The relatively divergent viral stimuli regulatory response is in part explained by a stronger response to GARD for the HG-Batwa individuals compared to their AG-Bakiga agriculturalist neighbors. Among the PopDR genes, the absolute fold-response to the viral ligand GARD was significantly stronger in the HG than the AG individuals (Figure 2C, Mann-Whitney-Wilcoxon Test *P* = 7.74×10^−32^), while a similar difference was not observed for LPS (Mann-Whitney-Wilcoxon Test; *P* = 0.34). Our data thus suggest that differences in viral exposure may have been a main factor contributing to the immune response divergence between the HG-Batwa and the AG-Bakiga.

While we do not have historical records of the viruses encountered by these populations, we can measure antiviral antibodies in present-day populations to gather information about their viral exposure. We used VirScan^18^ – a high-throughput method that allows comprehensive analysis of antiviral antibodies – to measure in all our samples serum antibodies against 130 viruses known to be present in Africa (see Materials and Methods). In measuring the relative variation of epitope burden found among the 130 viruses tested, we identified antibodies against 35 viruses (27%) whose levels were significantly different (FDR < 0.05) between HG and AG ancestry individuals (see Materials and Methods). Among these 35 viruses, 32 (91.4%) showed a higher burden (i.e., increased seropositivity) in individuals of HG-Batwa ancestry (Figure 2D). We observed increased seropositivity for only three viruses, all of which were human-specific single strand RNA viruses, in the AG individuals. Interestingly, viruses with higher burdens in the HG-Batwa population were significantly enriched for double stranded DNA viruses (20 of 32 observed; 14 of 31 expected; OR=3.7 (CI 1.5-9.9); Figure 2D; Fisher’s Exact test *P* =2.9×10^−3^), compatible with the hypothesis that DNA viruses are able to persist more readily in smaller populations than RNA viruses due to longer periods of latency ^19-21^. Though the differences reported herein may not be indicative of historical exposure, they do support the possibility that rainforest hunter-gather and agriculturalist populations (at least in southwest Uganda) have faced significant differences in viral exposure, with rainforest hunter-gatherer populations exhibiting a higher viral burden, particularly when considering DNA viruses.

### Genetic variation significantly contributes to ancestry-associated differences in immune regulation

Next, we aimed to identify components of the HG and AG transcriptional immune response driven by either genetic or environmental factors between HG and AG populations. To limit the effects of unknown confounding factors, we used a linear regression model that accounts for population structure and principal components of the expression data (see Materials and Methods). We first identified genetic variants from the ~10.5 million genotyped SNPs that are associated with differences in gene expression levels (i.e., eQTL) in our complete sample. We focused specifically on cis-eQTL, which we defined as SNPs located either within or flanking (±100 kb) the gene of interest. We identified a total of 3,941 genes (37.6% of all genes tested) that are associated with at least one cis-eQTL (FDR<0.05) in at least one condition. Consistent with previous findings ^22-25^, a large fraction of cis-eQTLs (14.7%) were observed only in stimulated samples (Figure 3A, Figure 3B for an example), highlighting the key importance of gene-environment interactions to the transcriptional regulation of innate immune responses.

We then tested whether PopDE and PopDR genes were more likely to be influenced by genetic variants than expected by chance. We found that PopDE and PopDR genes were significantly enriched among the set of genes associated with cis-eQTLs (>1.6x fold-enrichment; *P* < 1.0×10^−10^; Figure 3C). These results suggest that the differences in transcriptional responses to viral and bacterial stimuli identified in HG- and AG-ancestry individuals are driven, at least partly, by genetic regulatory variants. To explicitly quantify the minimum contribution of identified cis-eQTL to the transcriptional differences detected between populations, we used the following approach. First, we estimated in each condition the proportion of variance explained (PVE) by HG-ancestry among PopDE genes. Then, we re-calculated HG-ancestry PVE after regressing out the effect of the single cis-SNP for each gene that was most strongly associated with the target gene’s expression level (i.e. the SNP with the lowest FDR, regardless of significance level). The difference between HG-ancestry PVE values before and after regressing out the cis-eQTL effect (normalized by the original PVE value) quantifies the proportion of ancestry-associated effects on gene expression that stems from the strongest cis-associated variant. Hereafter we refer to this score as ∆PVE. Using this approach, we estimated that cis-regulatory variants explain, on average, ~34% of the PopDE signal in each condition (average ∆PVE = 36.7%, 37.5% and 34.2% among PopDE genes (FDR < 0.2) in control, GARD and LPS condition, respectively; Supplementary Figure S4). From this analysis, we identified a set of 475 PopDE genes across conditions for which a single cis-eQTL is enough to explain almost all ancestry effects on gene expression levels (∆PVE > 75%; FDR<0.1; hereafter referred to as high-∆PVE variants) on gene expression levels (Figure 3D).

### Positive selection has helped shape immune response differences

We next examined whether positive selection has contributed to the identified differences in immune response between the HG and AG populations. To do this we focused specifically on the set of 475 high-∆PVE variants, which represent a genetic substrate on which natural selection could potentially act to drive differences in immune response between the two population groups. Given that AG populations have recently shifted their mode of subsistence (i.e. from hunting and gathering to agriculture), they are hypothesized to have experienced commensurate changes in pathogen burden and novel selection pressures^1-4^. Under this scenario, we would expect to observe stronger evidence of positive selection on high-∆PVE SNPs in the AG-Bakiga population relative to that observed for the HG-Batwa population. Surprisingly, our data suggest the opposite.

We found that high-∆PVE SNPs were significantly more likely to have extreme levels of population differentiation (i.e., Fst value above the 95^th^ percentile of the genome-wide distribution) as compared to equally-sized sets of SNPs matched for allele frequencies with high-∆PVE SNPs (Figure 4A, > 3.4-fold enrichment in all conditions; P. value < 10^−4^). This result suggests a driving role for evolutionary processes in shaping HG-Batwa and AG-Bakiga population divergence in immune regulation but does not alone distinguish the population lineage(s) on which the selection occurred. We therefore also calculated the population branch statistic (PBS)^26^, which provides an estimate of the magnitude of allele frequency change for each SNP that occurred along each population lineage following divergence from a common ancestor (see Materials and Methods). Using this statistic, we found that the majority of the allele frequency divergence among high-∆PVE SNPs occurred along the HG-Batwa lineage (mean PBS HG-Batwa = 0.16; mean PBS AG-Bakiga = 0.04; Mann-Whitney T-test P = 1.2×10^−14^), and not in the lineage leading to the AG-Bakiga population (Figure 4B). Importantly, the relative difference in the branch length leading to the HG-Batwa lineage *vs* the AG-Bakiga lineage among high-∆PVE SNPs is significantly greater than that based on genome-wide expectations (4.0 vs 2.3 in average out of 100,000 sets of randomly sample sets of 475 SNPs matched for allele frequencies to high-∆PVE SNPs, P=2.5×10^−4^).

**Fig. 4.**
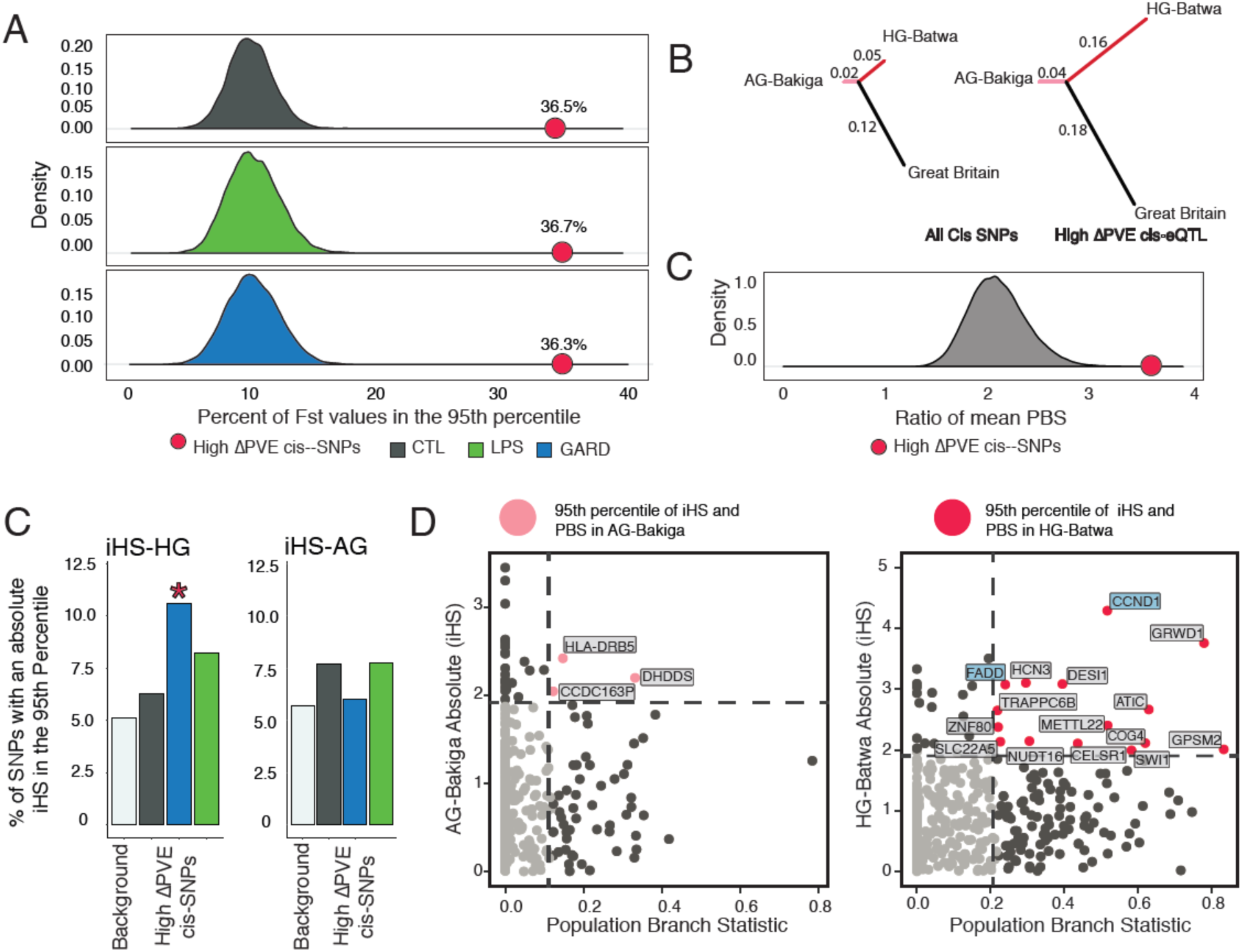
Evidence of selection driving population differences in immune response. (**A**) This density plot shows the distribution of the percent of SNPs with extreme values of Fst (e.g. in the 95^th^ percentile) for a set of randomly sampled cis-SNPs equally-sized sets of SNPs matched for allele frequencies with high-∆PVE SNPs. 10,000 iterations were run to obtain the distribution for each condition. The red point on each graph shows the percentage of high-∆PVE SNPs in the 95^th^ percentile. High-∆PVE variants in all conditions had significantly more SNPs in the 95^th^ Percentile (Fst comparison Chi-Squared Statistic; CTL P. value = 2.2^−16^, LPSP. value = 2.2^−16^, GARD P. value = 2.2^−16^). (**B**) A tree diagram illustrating the mean values of the population branch statistic for the HG-Batwa, AG-Bakiga, and a cohort from Great Britain as an outgroup. This figure illustrates a greater mean PBS score in the HG-Batwa population among high-∆PVE variants. (**C**) The distribution of the ratio of mean PBS in the HG-Batwa to the AG-Bakiga for a set of randomly sampled cis-SNPs equally-sized sets of SNPs matched for allele frequencies with high-∆PVE SNPs. 100,000 iterations were run to obtain the distribution and to calculate the P. value. The red point shows the ratio of mean PBS values represented as the branch lengths in the tree graph. (**D**) A bar graph illustrating the percentage of high-∆PVE SNPs that have an iHS value in the 95^th^ percentile compared to a background of all top cis-SNPs. For iHS, only values in the GARD-stimulated cells in the HG-Batwa population had significantly more SNPs in the 95^th^ percentile (HG-Batwa iHS comparison Chi-Squared Statistic; CTL P. value = 0.446, LPS P. value = 0.080, GARD P. value = 0.002; AG-Bakiga iHS comparison Chi-Squared Statistic; CTL P. value = 0.586, LPS P. value = 0.929, GARD P. value = 0.210). (**D**) PBS values for selection between populations graphed against absolute iHS values showing selection within each. population for high-∆PVE variants. Pink (AG-Bakiga) and Red (AG-Batwa) dots represent high-∆PVE SNPs in the 95^th^ percentile of both PBS and iHS. Among this group points are labeled with the corresponding gene name.

Additionally, we observed a significant enrichment of extreme integrated haplotype score (iHS) values (a neutrality test devised to detect recent positive selection events within a population)^27^ among high-∆PVE SNPs only in the HG-Batwa population. Specifically, we found that extreme iHS variants in the HG-Batwa population (>95^th^ percentile) were significantly enriched (2.1-fold) among high-∆PVE SNPs associated to GARD PopDE genes as compared to the set of all cis-SNPs (Chi-squared test, *P* = 1.75×10^−3^, Figure 4C). No such enrichments were observed in the AG-Bakiga population. Finally, more high-∆PVE SNPs and associated genes show strong signatures of natural selection (95^th^ percentile for both PBS and iHS) in the HG-Batwa (n=15) than in the AG-Bakiga (n=3) (Figure 4D), further supporting the conclusion that positive selection in the HG-Batwa lineage has at least partly led to the extreme levels of population differentiation observed in the set of high-PVE variants.

## Discussion

Our study provides the first genome-wide functional genomic comparison of variation in immune responses to infection between human hunter-gatherer and agricultural populations in Africa. Altogether, our results demonstrate that positive natural selection has contributed to present-day differences in innate immune responses between the HG-Batwa and the AG-Bakiga. Yet since functional evolutionary change occurred disproportionately on the HG-Batwa lineage, our results do not provide support for the long-standing hypothesis that selective pressures imposed by pathogens were particularly acute (at least in this region of the world) for agriculturalist populations due to the emergence of new crowd epidemic diseases. While it is difficult to contest the premise that the advent of agriculture led to the emergence of new pathogens and to the increased pathogenicity of others, it is likely that other, perhaps yet unknown, diseases have simultaneously been consistently more prevalent in hunter-gatherer populations. In particular, our serological data suggest that differences in viral exposure may have been a primary contributing factor to the divergence of HG-Batwa and AG-Bakiga immune responses. This notion is consistent with recent claims that viruses have been the primary drivers of adaptive evolution in mammals^28^ and one of the main selective pressures during recent human evolution^29^.

We chose to work with the HG-Batwa and AG-Bakiga for two reasons. First, while these two populations live in a relatively remote area of southwest Uganda, samples collected from this region could be transferred to a cell culture laboratory within 24 hours – a critical factor needed to ensure the viability of PBMCs – and processed identically, limiting possible batch effects that otherwise can affect inter-population functional comparisons. Second, while the long-term ecological histories of these two populations are distinct, they have shared similar environments and subsistence modes since 1992, when the HG-Batwa were evicted from Bwindi Impenetrable Forest. Thus, potential proximate environmental effects have been minimized to the greatest possible degree, facilitating our study of the genetic basis of functional genomic variation.

Yet, our study is still not free of challenges. First, our relatively small sample size – an inherent constraint when studying hunter-gatherer populations especially – limits our power to detect eQTL. Thus, it is likely that we are underestimating the true genetic contribution to ancestry-related differences in gene expression. Moreover, our ability to detect recent events of positive selection (such as those hypothesized to have occurred on immune system loci following the advent of agriculture) is bounded by the limited power of the currently available neutrality tests^27^, especially if selection occurred on standing genetic variation^30^. We also emphasize that these population lineages diverged more than 60,000 years ago, long prior to the origins of agriculture in Africa (refs). Thus, a substantial proportion of the functional genetic divergence we observed likely reflects earlier evolutionary responses to longstanding ecological differences facing each lineage. Still, our results are in direct opposition to *a priori* expectations of radical shifts in selection pressures on human immune systems following the agricultural transition, suggesting that the reality may instead be much less straightforward. Future studies on immune responses to a larger array of pathogenic stimuli, in additional cell types, and on additional pairs of hunter-gatherer and agriculturalist populations will help to more precisely characterize the impacts of agriculture on the evolution of human immune systems.

## Methods

### Sample collection

Blood samples were taken from a total of 103 individuals, 59 HG-Batwa (Hunter-gatherer) and 44 AG-Bakiga (Bantu speaking agriculturalist) individuals (see Supplementary Figure 1). We restricted our sample collection to adult individuals. For the HG-Batwa, we only collected samples from individuals who had lived in the forest and that were born prior to the 1991 formation of Bwindi Impenetrable Forest National Park, a time point known well to the HG-Batwa. The HG-Batwa and AG-Bakiga samples were collected under informed consent (Institutional Review Board protocols 2009-137 from Makerere University, Uganda, and 16986A from the University of Chicago). The project was also approved by the Uganda National Council for Science and Technology (HS617).

### Characterization of cell type composition

PBMCs were isolated from whole blood by Ficoll-Paque centrifugation and cryopreserved. Cell type composition of each PBMC sample was quantified using the following conjugated antibodies: CD3-FITC (clone UCHT1, BD Biosciences), CD20-PE (clone L27, BD Biosciences), CD8-APC (clone RPA-T8, BD Biosciences), and CD4-V450 (clone L200, BD Biosciences), CD16-PE (clone 3G8, Biolegend), CD56-APC (clone HCD-56), and CD14-Pacific Blue (clone M5E2, Biolegend). Antibodies were incubated for 20 min. Fluorescence was analyzed on a total of 30,000 cells for each population per sample with a FACSFortesa (BD Biosciences) and the FlowJo software (Treestar, Inc., San Carlos, CA). Supplementary Figure 2 illustrates what combinations of markers were used to define each of the cellular populations we considered in this study.

### Ligand stimulation

PBMCs were cultured in RPMI-1640 (Fisher) supplemented with 10% heat-inactivated FBS (FBS premium, US origin, Wisent) and 1% L-glutamine (Fisher). For each of the tested individuals, PBMCs (2 million per condition) were stimulated for 4 hours at 37°C with 5% CO_2_ with the immune challenges gardiquimod (GARD, 0.5µg/ml, TLR7 and TLR8 agonist) or lipopolysaccharide-EB (LPS, 0.25 µg/ml, TLR4 agonist). A control group of non-stimulated PBMCs were treated the same way but with only medium.

### Steps for RNA-Sequencing

Total RNA was extracted from the non-stimulated and stimulated cells using the miRNeasy kit (Qiagen). RNA quantity was evaluated spectrophotometrically, and the quality was assessed with the Agilent 2100 Bioanalyzer (Agilent Technologies). Only samples with no evidence of RNA degradation (RNA integrity number > 8) were kept for further experiments. RNA-sequencing libraries were prepared using the Illumina TruSeq protocol. Once prepared, indexed cDNA libraries were pooled (6 libraries per pool) in equimolar amounts and sequenced with single-end 100bp reads on an Illumina HiSeq2500. In total we generated RNA-sequencing profiles for 265 samples coming from 101 different individuals.

Adaptor sequences and low-quality score bases (Phred score < 20) were first trimmed using Trim Galore (version 0.2.7). The resulting reads were then mapped to the human genome reference sequence (Ensembl GRCh37 release 75) using STAR (2.4.1d)^**31**^ with an hg19 transcript annotation GTF downloaded from ENSEMBL (date: 2014-02-07). Reads matrices were computed using htseq-count^**31**^. To ensure stringent quality control of the RNA-seq data we removed from downstream analyses samples: (i) with less than 10 million of sequencing reads, (ii) with less than 50% of reads mapping to annotated exons; and (iii) samples that in a principal component analysis appeared to be contaminated or had failed to respond to the immune challenges. After these filtering steps we were left with 229 samples (76 CTL, 83 LPS and 70 GARD, samples labeled as PopDE_set=1 in Supplementary Figure 1), coming from 99 individuals (42 Bakiga, 57 Batwa).

### Identification of PopDE genes

To estimate the effects of HG ancestry on gene expression (within each experimental condition), gene expression levels across samples were normalized using the TMM algorithm (i.e., weighted trimmed mean of M-values), implemented in the edgeR R package^32^. Afterwards, we log-transformed the data and obtained precision-weights using the voom function in the limma package^**33**^. Only genes showing a median log2(cpm) > 2 within at least one of the experimental conditions were included in the analyses, which resulted in a total of 10,895 genes. Sequencing Flowcell batch effects were removed using the function ComBat, in the sva Bioconductor package^34^. Then, expression was modelled as a function of hunter-gatherer ancestry (HG) levels, while correcting for sex (x_1_), proportions of CD4^+^ T-cells (x_2_), CD14^+^ monocytes (x_3_), CD20^+^ B-cells (x_4_) and the fraction of reads assigned to the transcriptome (x_5_). Monocytes, T-cells and B-cells were included in the model after we identified that they were the only significant drivers of tissue composition effects on gene expression (cell types whose proportion in blood had a significant impact (FDR<5%) in at least 2.5% of the genes tested, in at least one condition).

Using the weighted fit function from limma (lmFit) and the weights obtained from voom, we fitted the following model:

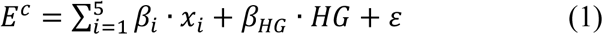

Where *E*^*c*^ represents the vector of flowcell-corrected expression levels of a given gene in condition *c*, *β*_*i*_ the effects of the covariates, and *β*_*HG*_ the effect of hunter-gatherer genetic ancestry. The *β* of these coefficients represent the fold-change (FC) effects associated to unit variation in each of the variables tested. This means, for sex, the average differences in expression between male and female, for HG, (FC) between HG and AG, while, the rest of the variables, since they are standardized, represent the differences in expression associated to a shift in the covariate equal to one standard deviation.

### Estimation of PopDR statistics

In order to model the effects of HG admixture on the intensity of the response to either GARD or LPS stimulation (i.e. PopDR effects), individual-wise fold-changes matrixes were built for each ligand. To do so, the effects of the technical covariates (i.e. sex, tissue composition and fraction of mapped reads) were first removed from the Flowcell-corrected expression matrixes within each condition. The resulting matrixes were subtracted (i.e. LPS-CTL and GARD-CTL, in log2 scale) to build corrected fold change matrixes using for that end only individuals for which pairs of samples CTL vs ligand were available (70 individuals for LPS, 59 for GARD, see Supplementary Figure 1). Finally, fold-changes were modeled according to a simple design *FC* = *β*_HG_ · *HG* + ε, using lmFit, with weights propagated from the ones calculated by voom for each condition. More specifically, voom weights are the inverse of the variance expectation for each RNAseq entry, obtained from the method defined by Robinson et al.^33^. That means that, if, for a given fold-change entry *FC* = *E*^*ligand*^ − *E*_*CTL*_, we propagate the expected variance of the FC as follows: *σ*^2^(*FC*) = *σ*^2^(*E*^*ligand*^) + *σ*^2^(*E*^*CTL*^).

Since the within condition weights were: *w*_*ligand*_ = 1/*σ*^2^(*E*^*ligand*^) and *w*_*CTL*_ = 1/*σ*^2^(*E*^*CTL*^), *σ*^2^(*FC*) = 1/*σ*^2^(*E*^*ligand*^) + 1/*σ*^2^(*E*^*CTL*^), and, finally:

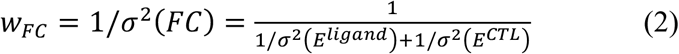

### Ligand stimulation effects and DE statistics

In order to estimate the overall LPS and GARD effects on gene expression, we separated the samples as CTL+GARD and CTL+LPS samples and analyzed them following the same analytical procedure used for PopDE, this time according to the following model design:

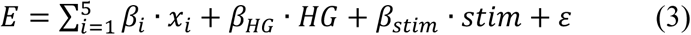

where *stim* is a dummy variable capturing the association of each sample to either the CTL condition (stim=0), or the stimulated condition (stim=1), and, thus, *β*_*stim*_ captures the overall ligand effects on gene expression. Whilst the CTL and LPS samples were sequenced together as part of the same sequencing batch, the GARD samples were sequenced in a later batch. Thus, to avoid the confounding sequencing batch and the effects of GARD-stimulation, we re-sequenced a reduced number of CTL samples along with the GARD batch, of which, 5 CTL-samples passed our QC filters. We used these samples to obtain the GARD effects as described in this section (sample set labeled as GARD_stim_set=1 in Supplementary Table 1).

### False discovery rates in PopDE, PopDR, and stimDE analyses

To avoid biases related to distributional assumptions on statistical significance that might arise as a result of batch removal procedures or data pre-treatment, for all of our PopDE and popDR analyses, we controlled for multiple testing using a generalization of the false discovery rate method of Storey and Tibshirani, re-calibrated to empirical null p-value distributions generated via permutation tests, as we previously described^24^. To perform these tests, in the case of PopDE and PopDR effects, HG-Batwa admixture was randomly permuted, while for stablishing the null distribution for ligand stimulation effects, condition labels (CTL vs stimulus) were randomly re-assigned within each individual. In this case, whenever one single sample was available for a given individual, it was labeled either as CTL or stimuli (either LPS or GARD), with probability=0.5. Permutation tests were repeated 1000 times per test.

### Gene ontology enrichments

To identify functional enrichments among genes that were both significantly upregulated by the ligands and show differences in expression between populations in the stimulated conditions, we used the cytoscape app ClueGO (vesion 2.3.3)^35^. Specifically, we tested the enrichments of all GO terms between GO levels 4 and 7, using a Fisher-exact test. We corrected for multiple testing using the Benjamini-Hochberg method. To select a set of most relevant GO terms suitable for visualization, we selected, in figure 1F, terms with enrichment p<5×10^−4^ and either number of genes in target gene sets >8 or percentage of total term size in target >10, in at least one of the four columns represented. The complete results of these analyses are shown in Supplementary Table 3.

### Antibody profiling

Antibody profiling was performed using VirScan, as previously described^18^. Briefly, we added 2µl of sera to 1 ml of the VirScan bacteriophage library, diluted to ~2 × 10^5 fold representation (2 × 10^10^ plaque-forming units for a library of 10^5^ clones) in phage extraction buffer (20 mM Tris-HCl, pH 8.0, 100 mM NaCl, 6 mM MgSO4), in a single well of a 96-deep-well plate, pre-blocked with 3% bovine serum albumin in TBST. We allowed the serum antibodies to bind the phage overnight on a rotator at 4°C. To each well, we then added 40 µl of a 1:1 mixture of magnetic protein A:protein G Dynabeads (Invitrogen) and rotated for 4 hours at 4°C to allow sufficient binding of phage-bound antibodies to magnetic beads. Using a 96-well magnetic stand to immobilize the magnetic bead-antibody-phage complexes, we then washed the beads three times with 400 ml of PhIP-Seq wash buffer (50 mM Tris-HCl, pH 7.5, 150 mM NaCl, 0.1% NP-40). After the final wash, beads were re-suspended in 40 ml of water and phage were lysed at 95°C for 10 minutes. For downstream statistical analyses, we also lysed phage from the library before immunoprecipitation (the input library) and after immunoprecipitation using only phage extract buffer without serum ("beads only control"). Each sample was run in duplicate.

Briefly, we performed two rounds of PCR amplification on the lysed phage material using hot start Q5 polymerase. The first round of PCR used the primers IS7_HsORF5_2 and IS8_HsORF3_2. The second round of PCR used 1 ml of the first-round product and the primers IS4_HsORF5_2 and a unique indexing primer for each sample to be multiplexed for sequencing, where “xxxxxxx” denotes a unique 7-nt indexing sequence (See below). After the second round of PCR, DNA concentration was quantified using qPCR, and pooled equimolar amounts of all samples were used for gel extraction. The extracted pooled DNA was sequenced by the Harvard Medical School Biopolymers Facility using a 50– base pair read cycle on an Illumina HiSeq 2000 or 2500, with the full pool split and run over both lanes of a HiSeq flow cell to obtain 700,000 - 1,300,000 reads per sample.

IS7_HsORF5_2:

ACACTCTTTCCCTACACGACTCCAGTCAGGTGTGATGCTC

IS8_HsORF3_2:

GTGACTGGAGTTCAGACGTGTGCTCTTCCGATCCGAGCTTATCGTCGTCATCC

IS4_HsORF5_2:

AATGATACGGCGACCACCGAGATCTACACTCTTTCCCTACACGACTCCAGT

Indexing Primer:

CAAGCAGAAGACGGCATACGAGATxxxxxxxGTGACTGGAGTTCAGACGTGT

After sequencing, samples were deconvoluted and reads aligned to the known epitope reference library for quantification and statistical analysis, as previously described. When an antibody against a particular epitope was in the sample serum, the epitope was expected to be enriched above a specific threshold, with the threshold dependent on the relative input count of the particular phage in the input library. P-values for enrichment were calculated using generalized Poisson regression to obtain a distribution of NGS read counts per sample for a given input count.

### Analysis of viral epitope burden

The goal of this analysis was to identify viruses differentially associated to either one of the two populations tested. To that end, we first restricted our analysis to a set of 130 viruses known to be present in Africa. The full list of viruses tested can be found in Supplementary Table 4. For these viruses, we obtained an estimation of seropositivity for each individual by counting the number of epitopes for which they tested positive (defined as epitopes detected above background at a p<0.05 in both technical replicates). After filtering out lowly represented viruses (i.e. those whose median number of epitopes across all individuals was lower than 2), the number of viruses was reduced to 112, for which we quantified the relative deviation of epitope counts per individual, with respect to the overall mean of each virus. Explicitly, let 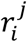 represent the number of positive epitopes for virus i and individual j, and 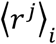 the virus average across all individuals. Thus, the relative deviation in seropositivity for each individual gets defined, for individual i and virus j as 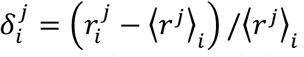. By testing for a linear association between 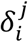 and HG ancestry, we estimate the inter-population differences in seropositivity relative to the mean epitope prevalence of each virus. We conducted this analysis using the lmFit function, in the R package limma^33^. Finally, false discovery rates associated to these linear models were estimated using Storey and Tibshirani’s method implemented in the R package qvalue^36^.

### Genome-wide genotyping

From the 99 individuals that were included in the sample-set used for PopDE analyses, a subset of 96 individuals (54-Batwa and 42-Bakiga, samples labelled as EQTL_set=1 in Supplementary Table S1) were successfully genotyped on the Illumina HumanOmni1-Quad genotyping array (Illumina, San Diego, USA), as previously described^16^. We got phased genotypes using ShapeIT v2 and obtained the imputed dataset using Impute2^37^. In total, 10,524,770 autosomal and X-linked SNPs passed quality control filters.

### Admixture and relatedness estimations

Admixture was estimated using a nonhierarchical clustering analysis of the SNP data using the software ADMIXTURE^17^, based upon independent SNPs (LD >0.3) from the genotyping chip dataset for the set of 96 individuals that were successfully genotyped. For the three individuals for which genotype data was not available (T15, T30 and T62, included in Pop_DE set but absent from EQTL_set), admixture values were estimated from the RNA-seq data. A pair-wise relatedness matrix among genotyped individuals was computed using Plink^38^. As expected, we found that the mean relatedness within each population was modest in both cases, but significantly larger among HG-Batwa (Mean relatedness among HG-Batwa samples: 6.9%; 0.6% among AG-Bakiga). To ensure that our results were not impacted by the increased number of related individuals in the HG-Batwa population, we re-ran our PopDE analyses excluding strongly related individuals (i.e., pi-hat > 0.375). This yielded 57, 58 and 62 samples in CTL, GARD and LPS condition, respectively (18, 12 and 21 samples removed in each condition, either because high relatedness or absent genotypes, of which 17, 10 and 20 were Batwa). The results of the PopDE analyses remained largely unaffected by the removal of these related samples (r > 0.94 for the correlation of the estimated effect sizes when using all the samples *vs* those obtained when we excluded closely related individuals; Figure S5).

### Mapping of cis – eQTL

Cis-eQTL mapping was conducted using the R package Matrix eQTL^39^. We estimated associations between SNP genotypes and changes in gene expression levels using a linear regression model where alleles affecting expression, denoted *G*, were assumed to be additive. This was conducted for each of the conditions separately with individuals from both populations included in the analyses to increase the power to map cis-eQTL. Associations of SNPs within the gene body or 100Kb upstream and downstream of the transcript start site and transcript end site were used to map cis-eQTL. SNPs with a minor allele frequency (MAF) less that 10% were removed from the analyses resulting in 2,284,380 autosomal SNPs that were tested against a total of 10,479 protein coding genes. To account for false positives resulting from population structure, the first two principal components obtained from a PCA on the genotype data were included in the model (*GPC*). For each library, we also took into account the potential biases and significant technical confounders. These included, as in the DE analyses, sex (x_1_), proportions of CD4^+^ cells (x_2_), CD14^+^ cells (x_3_), CD20^+^ cells (x_4_), the fraction assigned e.g the percentage of reads mapping to the transcriptome (x_5_), as well as sequencing flowcell, which was accounted for by including in the model as many covariates as sequencing flowcell levels *sf*_*i*_ present in each case (*n*_*sf*_(*c*)):

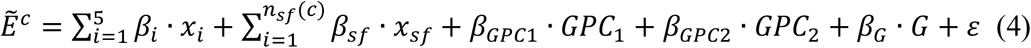

In this model, 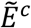 represents a vector of transformed expression values in condition c, which we obtained from the original expression values *E*_*c*_ after accounting for unmeasured-surrogate confounders. Specifically, we extracted the principal components *EPC*_*i*_ from a correlation matrix of the expression table within each condition *E*^*c*^, and then regressed out the first *n*_EPC_(*c*) of them as follows: 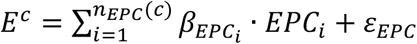; in order to obtain from the residuals of this expression the transformed expression values used in eq. (4): 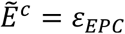. The specific number of PCs to regress out for each condition was chosen empirically (*23,25*), upon optimization of the signal strength obtained for EQTLs in eq. 4. This yielded *n*_*EPC*_(CTL) = *n*_*EPC*_(*GARD*) = 8 and *n*_*EPC*_(LPS) = 11.

### Proportion of Variance (PVE) estimations

In order to compute the proportion of variance explained (*PVE*) by the different covariates in the PopDE models (Supplementary Figure 3), we used the method proposed by Shabalin et al^40^, and implemented in the R package relaimpo^41^. According to this approach, the contribution of each covariate to the overall determination coefficient *R*^2^ is calculated upon adding sequentially all covariates to the model and calculating their contribution to the increase of *R*^2^ in each case, averaging across all possible covariate orderings. We summed the contributions of the three fractions of cell types included in the models (CD14^+^, CD4^+^ and CD20^+^) to obtain the estimates of tissue composition reported in the Supplementary Figure 3. The PVE associated either to sex (*PVE*_*sex*_), tissue composition (*PVE*_*tissue*_ = *PVE*_CD4_ + *PVE*_*CD4*_ + *PVE*_*CD20*_) and Hunter-gatherer ancestry (*PVE*_*HG*_), add up to the total fraction of explained variance for each gene, that is:

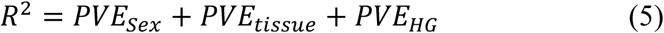

To quantify what fraction of the inter-population differences in gene expression was accounted for by cis eQTL, we first estimated, for each gene, the contribution of HG ancestry on gene expression variation within each condition (i.e. the PopDE effect-sizes 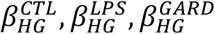, for genes showing statistical evidence of ancestry effects at a relaxed threshold of FDR<0.2). The proportion of variance explained by Hunter-gatherer ancestry 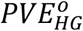 is defined as the increase in variance explained (that is the increase in *R*^2^) by the PopDE model in eq. 1, upon adding the HG variable as the last co-variable. Then, we fitted an alternative PopDE model for each gene, starting from equation (1), but adding the genotype of the top cis-SNP for the gene being tested, *G*_*Top*_, as follows:

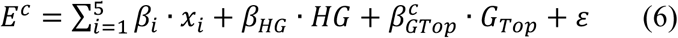

From this model, an analogous estimate 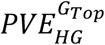 was obtained, which captured the relevance, in terms of explained variance, of adding hunter-gatherer ancestry, once the best SNP was already included in the model.

Once the contribution to final variance explained was obtained from both models we retrieved the difference between the two models 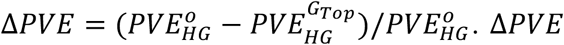 represents the proportion of the population difference in gene expression that can be attributed to the strongest cis eQTL for the gene of interest.

To assess the statistical significance of ∆*PVE*, we used the same approach described above but we removed the effect of the strongest cis-eQTL identified after randomly shuffling individual labels from the genotype data. Then, to construct a null model that was unbiased by the selection of the best SNP per gene, we built a third linear model, analogous to that of eq. (6) using, instead of the true, most significant SNP variant for that gene **G**_***Top***_, the most significant variant that arises by chance, among all the permuted SNPs: 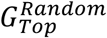:

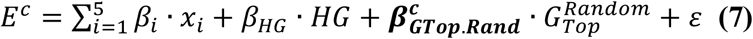

Then, we calculate PVE values based on the HG-admixture effects inferred from eq. 7, which we call 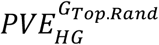. Finally, we estimate the null-expectation for ∆*PVE*, which we call ∆*PVE*_*null*_, as follows:

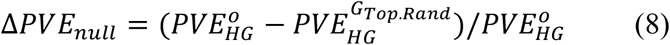

Comparing the distribution of observed ∆*PVE* to the distribution of its empiric null expectation ∆*PVE*_*null*_ we obtain empiric one-tailed p-values for each test, defined as the fraction of null-tests with ∆*PVE*_*null*_ > ∆*PVE*. Finally, proper correction for multiple testing (Storey-Tibshirani FDRs) of these empiric p-values allows us to stablish an empiric model for statistical significance of these effects (see Supplementary Figure 4).

### Selection Statistics

We calculated the selection statistics by using the individuals used to map cis-eQTL that had an admixture less than 0.2 or greater than 0.8 to clearly define the two populations. This included 43 Bakiga individuals and 39 HG-Batwa individuals. We calculated the fixation indexes (Fst) using a modified version of Wright’s Fst for all SNPs using VCFtools v0.1.12b^42^. The integrated haplotype scores (iHS) were calculated using Selscan, which is a program that calculates haplotype-based scans for recent or ongoing signatures of positive selection. This method is based on the knowledge that when adaptive de novo mutations quickly increase in frequency it reduces genetic diversity around this variant faster than recombination can occur. Therefore, this score is a measure of haplotype homozygosity extending from an adaptive locus^43^. To do this, phased genotypes were created using SHAPEITv2^44^ for each chromosome independently. We calculated iHS separately for the HG and AG population for all imputed genotypes. When estimating mean Fst and iHS among cis-eQTL we combined cis-eQTL mapped in all conditions and selected the variant with the lowest P. value for a given gene resulting in one cis-SNP per gene. The Fst and/or iHS for that SNP was then considered in this analysis. Finally, the population branch statistic (PBS) was calculated from Fst values using a cohort from Great Britain available from the 1000 Genomes Project as an outgroup. Fst was first used to calculate population divergence as [T= -log(1-Fst)], and then PBS was calculated for each SNP for HG-Batwa and AG-Bakiga as:

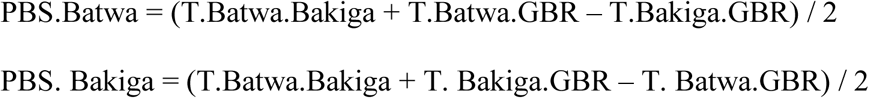

## Supporting information

## Acknowledgments

The authors would like to thank the Batwa and Bakiga communities and all individuals who participated in this study, and the Batwa Development Program, Byaruhanga Julius, Magambo Michael, Byamugisha Patrick, Twesigomwe Sabastian, Safari Joseph, and Busingye Levi for expert assistance during the sample collection process in Uganda. We also thank Nanyunja Sarah for technical laboratory assistance. We thank Jenny Tung and L.B.B. lab members for critical reading of the manuscript. We thank Calcul Québec and Compute Canada for providing access to the supercomputer Briaree from the University of Montreal. This work was supported by NIH R01-GM115656 to G.H.P and L.B.B., a fellowship from the Réseau de Médecine Génétique Appliquée (RMGA), and the Fonds de Recherche du Québec - Santé (FRQS) to G.F.H, and 1 F32 GM125228-638 01A1 to C.M.B. RNA-seq data have been deposited in Gene Expression Omnibus (accession number GSE120502). The 1M SNP genotype data are available at the European Genome-Phenome archive, www.ebi.ac.uk/ega/ (accession numbers EGAS00001000605, and EGAS00001000908).

## References

1 Diamond, J. & Bellwood, P. Farmers and their languages: the first expansions. Science 300, 597–603 (2003).

2 Greger, M. The human/animal interface: emergence and resurgence of zoonotic infectious diseases. Critical reviews in microbiology 33, 243–299 (2007).

3 Pearce-Duvet, J. M. The origin of human pathogens: evaluating the role of agriculture and domestic animals in the evolution of human disease. Biol Rev Camb Philos Soc 81, 369–382, doi:10.1017/S1464793106007020 (2006).

4 Wolfe, N. D., Dunavan, C. P. & Diamond, J. Origins of major human infectious diseases. Nature 447, 279–283, doi:10.1038/nature05775 (2007).

5 Gignoux, C. R., Henn, B. M. & Mountain, J. L. Rapid, global demographic expansions after the origins of agriculture. Proceedings of the National Academy of Sciences 108, 6044–6049 (2011).

6 Page, A. E. et al. Reproductive trade-offs in extant hunter-gatherers suggest adaptive mechanism for the Neolithic expansion. Proceedings of the National Academy of Sciences, 201524031 (2016).

7 Black, F. L. Measles endemicity in insular populations: critical community size and its evolutionary implication. Journal of Theoretical Biology 11, 207–211 (1966).

8 Anderson, R. M. & May, R. M. Infectious diseases of humans: dynamics and control. (Oxford university press, 1992).

9 Furuse, Y., Suzuki, A. & Oshitani, H. Origin of measles virus: divergence from rinderpest virus between the 11 th and 12 th centuries. Virology journal 7, 52 (2010).

10 Matthijnssens, J. et al. Full genome-based classification of rotaviruses reveals a common origin between human Wa-Like and porcine rotavirus strains and human DS-1-like and bovine rotavirus strains. Journal of virology 82, 3204–3219 (2008).

11 Suzuki, Y. & Nei, M. Origin and evolution of influenza virus hemagglutinin genes. Molecular biology and evolution 19, 501–509 (2002).

12 Sundararaman, S. A. et al. Genomes of cryptic chimpanzee Plasmodium species reveal key evolutionary events leading to human malaria. Nature communications 7, 11078 (2016).

13 Otto, T. D. et al. Genomes of all known members of a Plasmodium subgenus reveal paths to virulent human malaria. Nature microbiology 3, 687 (2018).

14 Barreiro, L. B. & Quintana-Murci, L. From evolutionary genetics to human immunology: how selection shapes host defence genes. Nature Reviews Genetics 11, 17 (2010).

15 Karlsson, E. K., Kwiatkowski, D. P. & Sabeti, P. C. Natural selection and infectious disease in human populations. Nature Reviews Genetics 15, 379 (2014).

16 Perry, G. H. et al. Adaptive, convergent origins of the pygmy phenotype in African rainforest hunter-gatherers. Proceedings of the National Academy of Sciences 111, E3596–E3603 (2014).

17 Alexander, D. H., Novembre, J. & Lange, K. Fast model-based estimation of ancestry in unrelated individuals. Genome research (2009).

18 Xu, G. J. et al. Comprehensive serological profiling of human populations using a synthetic human virome. Science 348, aaa0698 (2015).

19 McGeoch, D. & Davison, A. J. in Origin and evolution of viruses 441–465 (Elsevier, 1999).

20 McGeoch, D. J., Dolan, A. & Ralph, A. C. Toward a comprehensive phylogeny for mammalian and avian herpesviruses. Journal of virology 74, 10401–10406 (2000).

21 Van Blerkom, L. M. Role of viruses in human evolution. American Journal of Physical Anthropology: The Official Publication of the American Association of Physical Anthropologists 122, 14–46 (2003).

22 Barreiro, L. B. et al. Deciphering the genetic architecture of variation in the immune response to Mycobacterium tuberculosis infection. Proceedings of the National Academy of Sciences 109, 1204–1209 (2012).

23 Fairfax, B. P. et al. Innate immune activity conditions the effect of regulatory variants upon monocyte gene expression. Science 343, 1246949 (2014).

24 Nédélec, Y. et al. Genetic ancestry and natural selection drive population differences in immune responses to pathogens. Cell 167, 657–669. e621 (2016).

25 Quach, H. et al. Genetic adaptation and neandertal admixture shaped the immune system of human populations. Cell 167, 643–656. e617 (2016).

26 Yi, X. et al. Sequencing of 50 human exomes reveals adaptation to high altitude. Science 329, 75–78 (2010).

27 Voight, B. F., Kudaravalli, S., Wen, X. & Pritchard, J. K. A map of recent positive selection in the human genome. PLoS biology 4, e72 (2006).

28 Enard, D., Cai, L., Gwennap, C. & Petrov, D. A. Viruses are a dominant driver of protein adaptation in mammals. Elife 5, e12469 (2016).

29 Enard, D. & Petrov, D. A. RNA viruses drove adaptive introgressions between Neanderthals and modern humans. bioRxiv, 120477 (2017).

30 Prezeworski, M., Coop, G. & Wall, J. D. The signature of positive selection on standing genetic variation. Evolution 59, 2312–2323 (2005).

31 Dobin, A. et al. STAR: ultrafast universal RNA-seq aligner. Bioinformatics 29, 15–21 (2013).

32 Anders, S. et al. Count-based differential expression analysis of RNA sequencing data using R and Bioconductor. Nature protocols 8, 1765–1786 (2013).

33 Robinson, M. D., McCarthy, D. J. & Smyth, G. K. edgeR: a Bioconductor package for differential expression analysis of digital gene expression data. Bioinformatics 26, 139–140 (2010).

34 Ritchie, M. E. et al. limma powers differential expression analyses for RNA-sequencing and microarray studies. Nucleic acids research 43, e47-e47 (2015).

35 Leek, J. T., Johnson, W. E., Parker, H. S., Jaffe, A. E. & Storey, J. D. The sva package for removing batch effects and other unwanted variation in high-throughput experiments. Bioinformatics 28, 882–883 (2012).

36 Bindea, G. et al. ClueGO: a Cytoscape plug-in to decipher functionally grouped gene ontology and pathway annotation networks. Bioinformatics 25, 1091–1093 (2009).

37 Storey, J. D. & Tibshirani, R. in Functional Genomics 149–157 (Springer, 2003).

38 Howie, B. & Marchini, J. Instructions for IMPUTE version 2. (2009).

39 Purcell, S. et al. PLINK: a tool set for whole-genome association and population-based linkage analyses. The American Journal of Human Genetics 81, 559–575 (2007).

40 Shabalin, A. A. Matrix eQTL: ultra fast eQTL analysis via large matrix operations. Bioinformatics 28, 1353–1358 (2012).

41 Lindeman, R. H., Merenda, P. F. & Gold, R. Z. Introduction to bivariate and multivariate analysis. (Scott, Foresman Glenview, IL, 1980).

42 Grömping, U. Relative importance for linear regression in R: the package relaimpo. Journal of statistical software 17, 1–27 (2006).

43 Jeffrey, C. Genome-wide association study and meta-analysis finds over 40 loci affect risk of type 1 diabetes. Nat Genet 41, 703–707 (2009).

44 Szpiech, Z. A. & Hernandez, R. D. selscan: an efficient multithreaded program to perform EHH-based scans for positive selection. Molecular biology and evolution 31, 2824–2827 (2014).

