## Supplementary figures and images for "Natural selection contributed to immunological differences between human hunter-gatherers and agriculturalists"

### Supplementary file 1

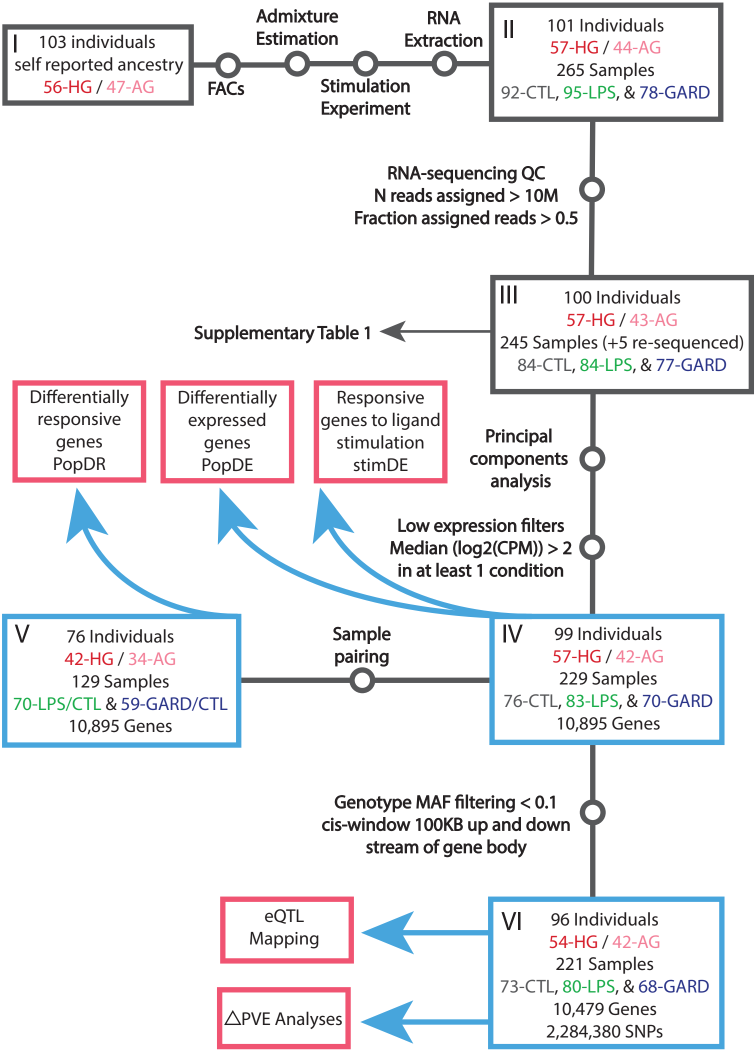

### Supplementary file 2

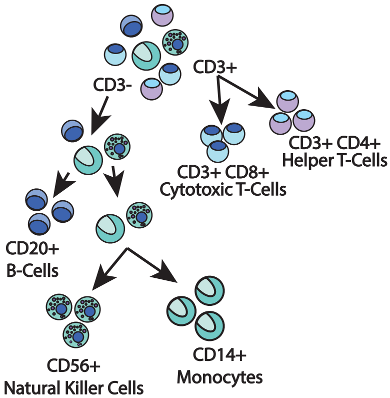

### Supplementary file 3

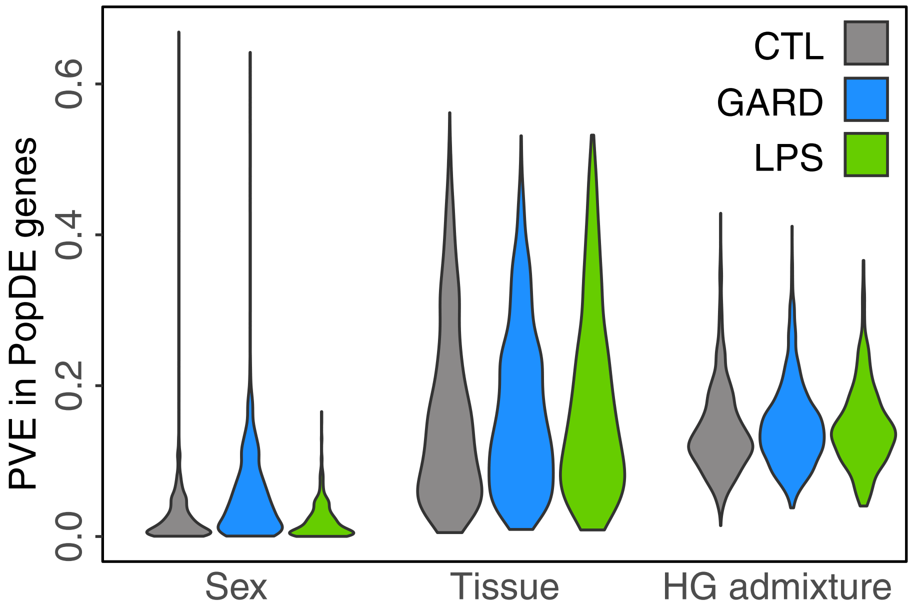

### Supplementary file 4

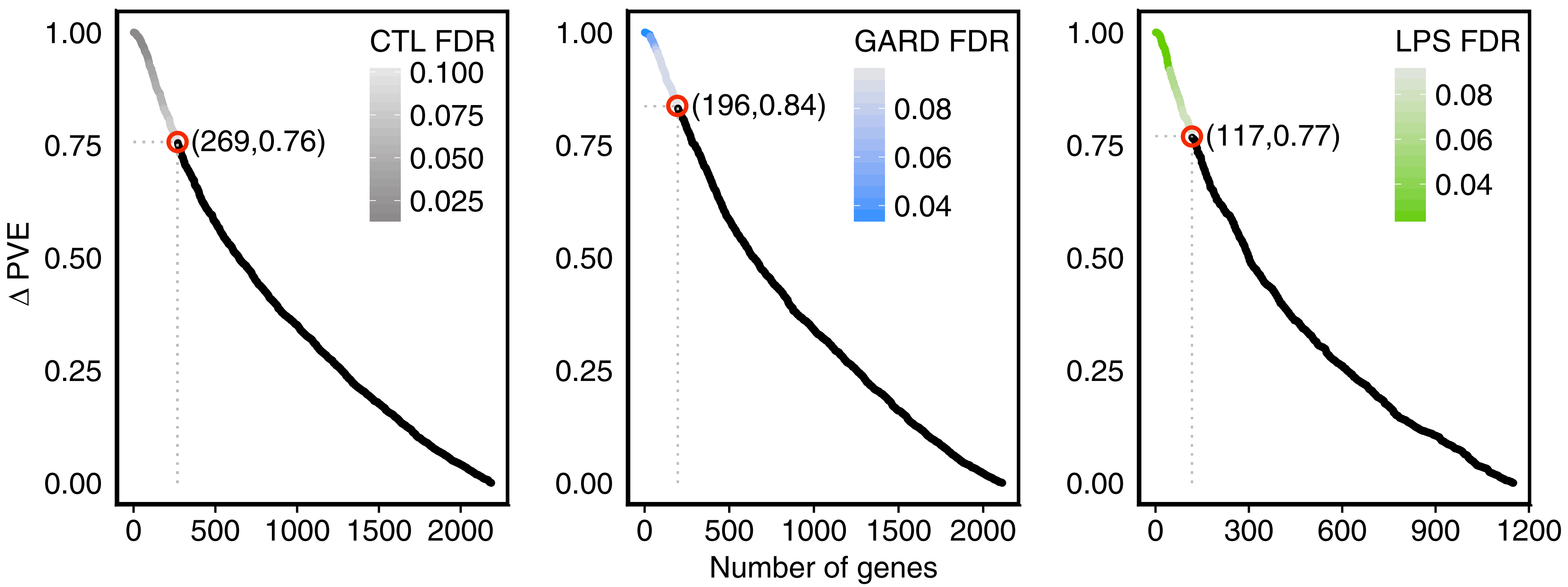

### Supplementary file 5

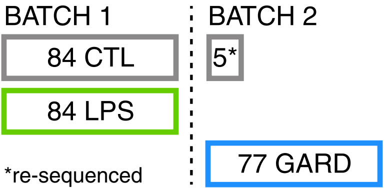

### Supplementary file 6

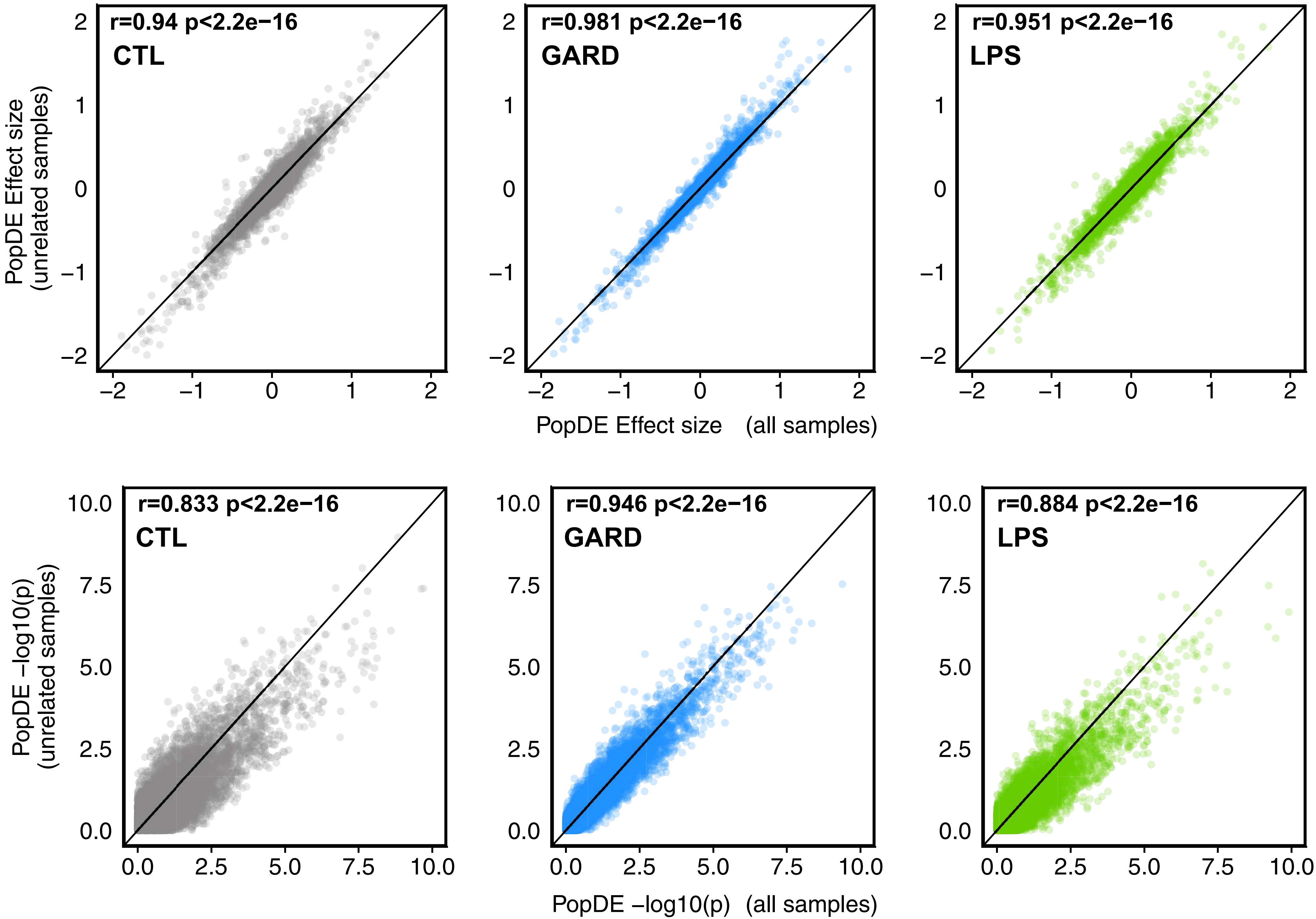

### Supplementary file 7

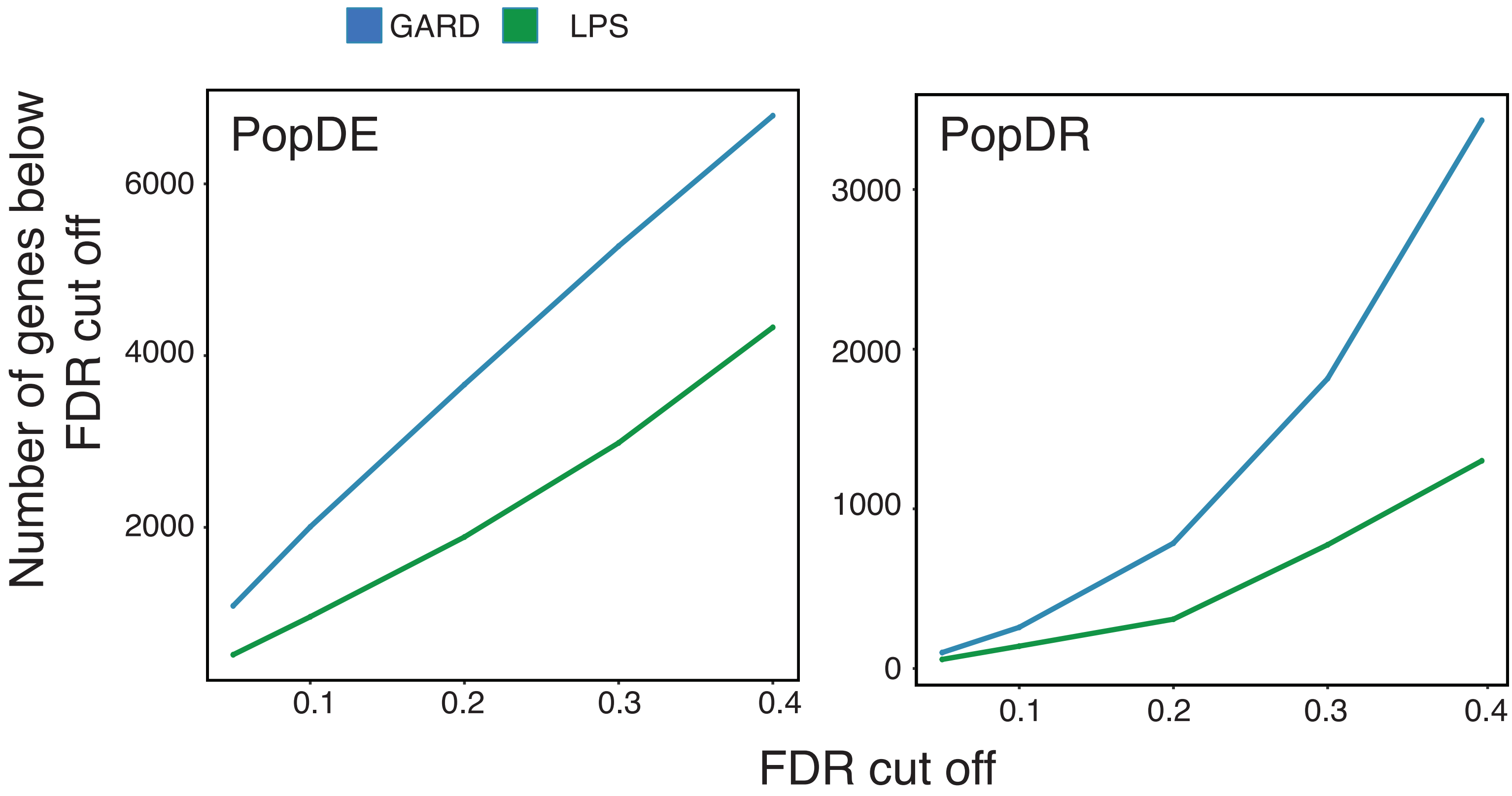
